# Enhancing AlphaFold-Multimer-based Protein Complex Structure Prediction with MULTICOM in CASP15

**DOI:** 10.1101/2023.05.16.541055

**Authors:** Jian Liu, Zhiye Guo, Tianqi Wu, Raj S. Roy, Farhan Quadir, Chen Chen, Jianlin Cheng

## Abstract

AlphaFold-Multimer has emerged as the state-of-the-art tool for predicting the quaternary structure of protein complexes (assemblies or multimers) since its release in 2021. To further enhance the AlphaFold-Multimer-based complex structure prediction, we developed a new quaternary structure prediction system (MULTICOM) to improve the input fed to AlphaFold-Multimer and evaluate and refine the outputs generated by AlphaFold2-Multimer. Specifically, MULTICOM samples diverse multiple sequence alignments (MSAs) and templates for AlphaFold-Multimer to generate structural models by using both traditional *sequence* alignments and new Foldseek-based *structure* alignments, ranks structural models through multiple complementary metrics, and refines the structural models via a Foldseek structure alignment-based refinement method. The MULTICOM system with different implementations was blindly tested in the assembly structure prediction in the 15th Critical Assessment of Techniques for Protein Structure Prediction (CASP15) in 2022 as both server and human predictors. Our server (MULTICOM_qa) ranked 3^rd^ among 26 CASP15 server predictors and our human predictor (MULTICOM_human) ranked 7^th^ among 87 CASP15 server and human predictors. The average TM-score of the first models predicted by MULTICOM_qa for CASP15 assembly targets is ∼0.76, 5.3% higher than ∼0.72 of the standard AlphaFold-Multimer. The average TM-score of the best of top 5 models predicted by MULTICOM_qa is ∼0.80, about 8% higher than ∼0.74 of the standard AlphaFold-Multimer. Moreover, the novel Foldseek Structure Alignment-based Model Generation (FSAMG) method based on AlphaFold-Multimer outperforms the widely used sequence alignment-based model generation. The source code of MULTICOM is available at: https://github.com/BioinfoMachineLearning/MULTICOM3.

## 1. Introduction

Single-chain proteins (monomers) often interact with each other to form multimers (i.e., assemblies or complexes) to perform functions such as gene regulation and signal transduction. The quaternary structures of multimers largely determine their function. Therefore, it is important to predict the quaternary structure of protein complexes from their sequences for studying protein-protein interaction and function. However, predicting the quaternary structure of protein complexes is more difficult than predicting the tertiary structure of single-chain monomers because the former involves more than one protein chain and needs to consider both intra-chain residue-residue interaction and inter-chain residue-residue interaction.

Prediction of protein complex structures has been traditionally carried out by molecular docking simulation techniques (e.g., simulated annealing or Markov Chain Monte Carlo simulation) guided by an energy or statistical potential function for decades^1, 2^. However, the accuracy of the protein docking is generally low^3, 4^. The recent application of deep learning to inter-protein contact prediction and quaternary structure prediction has started to transform the field^5–10^. Particularly, the adaption of the high-accuracy tertiary structure prediction method - AlphaFold2^11^ - for quaternary structure prediction as AlphaFold-Multimer^8^ has drastically improved the accuracy of quaternary structure prediction for protein assemblies.

Despite the breakthrough made by AlphaFold-Multimer, its accuracy for quaternary structure prediction is still much lower than AlphaFold2’s accuracy for tertiary structure prediction. Therefore, there is still a large room to further improve the accuracy of AlphaFold-Multimer-based complex structure prediction.

In this work, we developed several algorithms to improve AlphaFold-Multimer-based complex prediction from different aspects and integrated them to build a MULTICOM complex structure prediction system. It uses both traditional *sequence* alignments and new Foldseek^12^-based *structure* alignments to generate multiple sequence alignments (MSAs) for monomers and concatenates them as MSAs for multimers according to different criteria such as the same species and known/hypothetical protein-protein interaction. The structural templates identified by the sequence or structure alignments for monomers from different template databases are also combined as templates for the multimers. The diverse set of MSAs and templates are used as input for AlphaFold-Multimer to generate quaternary structural models, which are then ranked by multiple complementary model quality assessment methods including AlphaFold-Multimer’s confidence score, the average pairwise structural similarity (PSS) between a model and other models of the same target, and the average of the two. The top ranked models are further refined by using the Foldseek structure alignment-based model refinement to generate better models.

We implemented the MULTICOM system as two server predictors and two human predictors that blindly participated in the assembly structure prediction in CASP15 from May to August 2022. Both the MULTICOM server and human predictors ranked among the top server or human/server predictors in CASP15. The predictors also performed significantly better than a standard AlphaFold-Multimer predictor participating in CASP15, demonstrating that the MULTICOM approach has significantly improved the accuracy of the AlphaFold-Multimer-based protein assembly structure prediction. We released the source code of the MULTICOM system at GitHub so that the community can run it on top of AlphaFold-Multimer to obtain more accurate protein complex structure predictions.

## 2. Materials and Methods

### 2.1 The MULTICOM protein complex structure prediction system and methods

The workflow of the MULTICOM complex/multimer prediction system consists of seven steps (**Figure 1**): **(1)** *single-chain (monomer) structure prediction for each unit of a multimer*, **(2)** *multimer MSA generation*, **(3)** *multimer template identification*, **(4)** *multimer structural model generation*, **(5)** *Foldseek structure alignment-based multimer model generation*, **(6)** *multimer structural model ranking*, and **(7)** *Foldseek structure alignment-based refinement*. The method in each step is described as follows.

**Figure 1.**
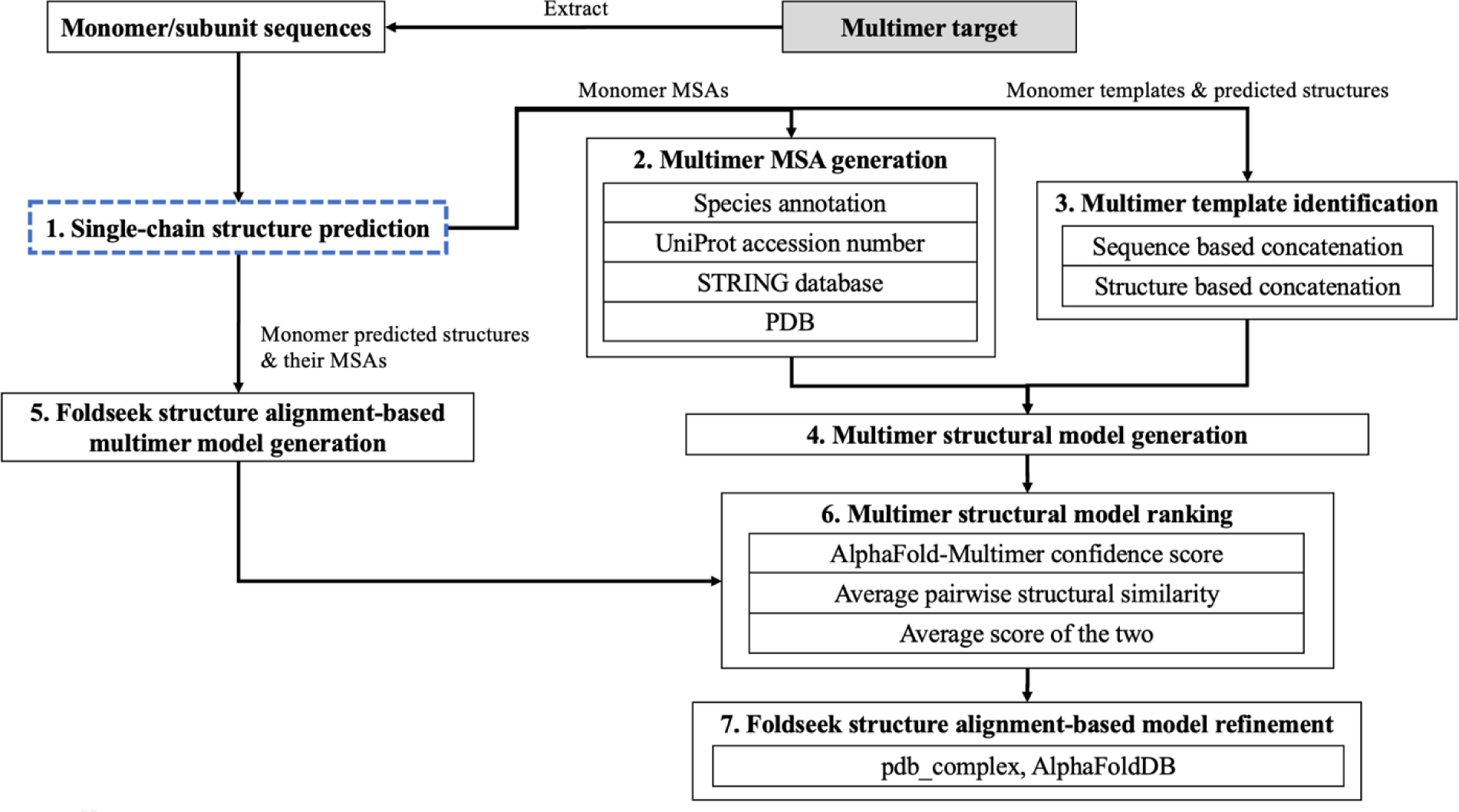
The workflow of the MULTICOM protein complex structure prediction system.

#### Single-chain structure prediction for each subunit of a multimer

Our in-house single-chain (monomer) tertiary structure prediction system^13^ built on top of AlphaFold2 is used to generate multiple sequence alignments (MSAs), structural templates, and predicted tertiary structures for each subunit of a multimer target. It uses sequence alignment tools including HHblits^14^, JackHMMER^15^, MMseq2^16^, an in-house implementation of DeepMSA^17^ to search multiple protein sequence databases including UniClust30^18^ (uniclust30_2018_08), UniRef30^18^ (UniRef30_2021_02), Uniref90^19^ (version 04/24/2022), UniProt^19^ (version 04/24/2022), the Integrated Microbial Genomes (IMG) database^20^ and the metagenome sequence databases (e.g., BFD^21, 22^, Metaclust^22^, MGnify clusters^23^) to generate a diverse set of MSAs for each unit (monomer).

#### Multimer MSA generation

AlphaFold-Multimer uses two kinds of MSAs as input: (1) the unpaired MSA for each subunit (MSA_unpaired_) and (2) the paired MSA that may encode the coevolutionary information between the subunits (MSA_paired_), which are prepared as follows by the MULTICOM system.

For hetero-multimers, the alignments in the MSAs of the subunits are concatenated using the potential protein-protein interaction information extracted from multiple sources to construct MSA_paired_ as shown in **Table 1**, including species annotations, UniProt accession number of sequences, protein-protein interactions in the STRING database^24^ and the complex structures in the Protein Data Bank^25^ (PDB). The alignment description (header) in UniClust30, UniRef30, UniRef90 and UniProt contains the UniProt ID, UniProt accession number and the species annotations (e.g., Organism identifier (OX)^8^, Organism name (OS)^26^, Taxonomy identifier (Tax)^26^). Based on the species information, the individual sequence alignments in the MSAs of the subunits belonging to the same species are concatenated to generate the paired multimer sequence alignments sequentially in a top-down manner. Based on UniProt accession numbers, sequence alignments from the subunit MSAs are concatenated if the difference between their UniProt accession numbers is smaller than 10 as in RosettaFold^27^. For simplification, the alignments with the same UniProt accession number prefix (e.g., except for the last character) are paired. The STRING database contains many hypothetical protein-protein interactions, each of which has an interaction score. The interaction score between two protein sequences in the UniProt database is retrieved according to the mapping between STRING ID to the UniProt ID. Two sequence alignments from two subunit MSAs are concatenated if their interaction score is higher than 500. According to the mapping between the PDB code and UniProt ID, two sequence alignments from two subunit MSAs are concatenated if they are mapped to the same PDB code indicating that they are two subunits of the same protein complex. The four sources of potential protein-protein interactions above are used by MULTICOM to generate 13 kinds of MSA_paired_ for hetero-multimers from the different databases (**Table 1**). The MSA_unpaired_ for hetero-multimers is always generated by the same default MSA generation procedure in AlphaFold2-Multimer (e.g., searching the subunit/chain sequence against UniRef30 and BFD, and MGnify clusters to generate MSAs). Both MSA_paired_ and MSA_unpaired_ are used in the model generation for hetero-multimers by AlphaFold-Multimer.

**Table 1.** The four sources of protein-protein interaction information used with four sequence databases for generating 13 kinds of paired MSAs for multimers in total.

| Sources of interaction | Interaction identifier | Sequence database | Number of kinds of paired MSAs |
| --- | --- | --- | --- |
| Species annotation | Organism identifier (OX),<br>Organism name (OS),<br>Taxonomy identifier (Tax) | UniClust30, UniRef30,<br>UniRef90, UniProt | 4 |
| UniProt accession number | Distance between UniProt accession numbers | UniRef30, UniRef90, UniProt | 3 |
| STRING database | Interaction score | UniRef30, UniRef90, UniProt | 3 |
| PDB complex | Same PDB code | UniRef30, UniRef90, UniProt | 3 |

For homo-multimers, the default MSA generation of the AlphaFold-Multimer is used by MULTICOM on different sequence databases to generate several kinds of MSA_paired_ (see *default_multimer*, *default_pdb*, *default_pdb70*, *default_comp*, *default_struct*, *default_af*, and *default_img* in **Table S1**) where the MSA_unpaired_ is simply concatenated horizontally together as the MSA_paired_ since the MSA_unpaired_ of each subunit is identical. In contrast, the customized multimer MSA generation methods in **Table S1** pair only the alignments in the MSAs of the subunits of the multimer that have the same species annotation or PDB complex codes to generate MSA_paired_. The alignments in the MSAs of the subunits without the species annotation or whose UniProt IDs cannot be mapped to any PDB codes are paired with gaps. Only MSA_paired_ is used in the model generation for homo-multimers by AlphaFold-Multimer, while MSA_unpaired_ is ignored.

#### Multimer template identification

The sequences of the subunits in the multimer are searched against the publicly available pdb_seqres database (version 04/24/2022), pdb70 (version 03/13/2022) monomer template database^11^ curated from Protein Data Bank (PDB), an in-house monomer template database pdb_sort90^13^, and an in-house protein template database (pdb_complex) constructed from only the biological assemblies in the PDB using HHSearch^28^, resulting four kinds of templates. pdb_complex was constructed in a similar way as pdb_sort90 except that the former only considered the biological assemblies in the PDB while pdb_sort90 considered all the proteins in the PDB. The templates found for each subunit are concatenated together if they share the same PDB code. Only one concatenated multimer template is kept for each PDB code. Finally, the predicted tertiary structures for each subunit/chain are also used as the fifth kind of templates, which can lead to highly inflated AlphaFold-Multimer confidence scores for generated models and is less useful than the other four kinds of templates (data not shown).

#### Multimer structural model generation

A customized version of AlphaFold-Multimer that accepts pre-generated MSAs (e.g., MSA_unpaired_ and MSA_paired_) and structural templates above as input is used to generate models. To perform more extensive model sampling, the value of parameter *num_ensemble* is changed from 1 to 3 and *num_cycle* from 3 to 5 in the customized AlphaFold-Multimer. The customized AlphaFold-Multimer takes up to 19 combinations of MSAs and structural templates (**Table S1**) for homo-multimer and up to 29 combinations of MSAs and structural templates (**Table S2**) for hetero-multimer as input to generate 10 structural models for each combination by setting the value of *num_multimer_predictions_per_model* to 2. Only the top 5 models ranked by the AlphaFold-Multimer confidence scores for each MSA-template combination are added into the structural model pool for the multimer, resulting in up to 95 (or 145) models generated for each homo-multimer (or hetero-multimer) target.

#### Foldseek structure alignment-based model generation

Different from using the *sequence alignment*-generated MSAs and templates above as input for AlphaFold-Multimer to generate models, we developed a novel Foldseek *structure alignment*-*based* multimer model generation (FSAMG) method (**Figure 2**) to generate up to 25 models as follows. The predicted tertiary structures of the subunits of a multimer generated by AlphaFold2 are searched against both the pdb_complex template database and the tertiary structure models in the AlphaFoldDB (the version 1 released before March 2022) by a fast structure alignment tool - Foldseek - to identity similar structural hits. The output of the Foldseek search includes the e-value of the structural hits as well as the structural alignments between the target model and the hits, which are converted into the sequence alignments between them. The two sequence alignments for two subunits/chains of the multimer are paired if they come from the same PDB protein complex (i.e., sharing the same PDB code) or from the two non-overlapping domains of a hit in the AlphaFoldDB. Only one paired alignment is kept for each PDB code to avoid the redundancy in the paired alignments. The sequence alignments are added into the MSA_unpaired_ for each subunit if it is a hetero-multimer, while the paired sequence alignments are included into the MSA_paired_ for both hetero-multimer and homo-multimer. For hetero-multimers, MSA_unpaired_ is initialized as the MSA generated for each subunit by the tertiary structure prediction system, while MSA_paired_ is set empty initially. For homo-multimers, MSA_paired_ is initialized as the MSA generated by the tertiary structure prediction system. The similar structural hits of the subunits of the multimer in pdb_complex are concatenated as multimer templates if they share the same PDB code. The structure-alignment generated MSAs and the concatenated multimer templates are used as input for the customized AlphaFold-Multimer to generate 10 models. The top 5 models ranked by their confidence scores are added to the structural model pool for the multimer. This procedure is applied with 2-5 top-ranked tertiary structure models of the subunits of the multimer as described above to generate 10 to 25 models in total. This structure alignment-based method can find some similar structural hits for hard targets that sequence alignments methods cannot, leading to deeper MSAs and more structural templates, which can be used by AlphaFold-Multimer to generate better structure prediction.

**Figure 2.**
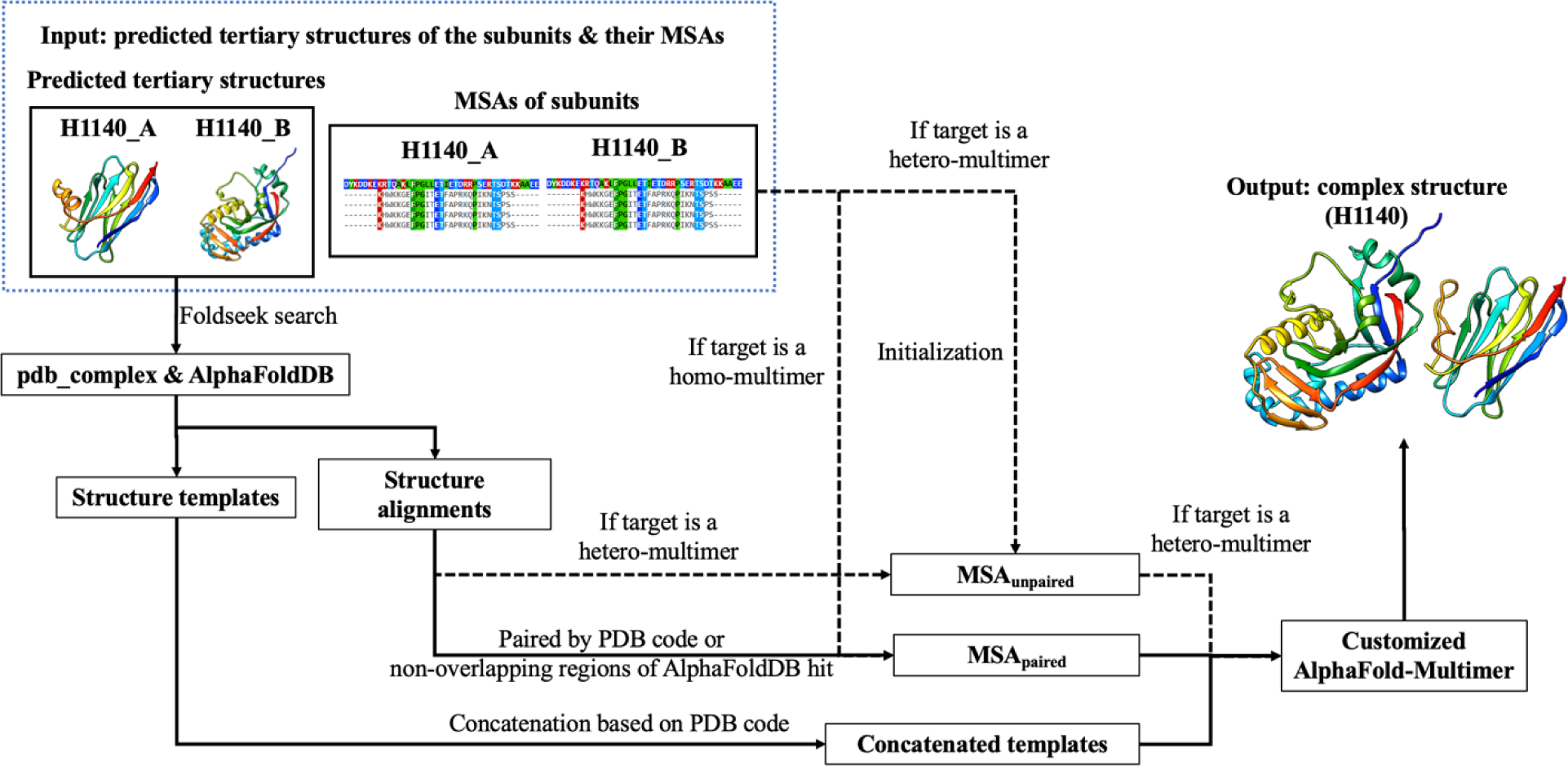
The illustration of the workflow of Foldseek structure alignment-based model generation (FSAMG) with a CASP15 hetero-dimer target H1140 as an example. For hetero-multimers, both MSA_unpaired_ and MSA_paired_ are used in model generation, while for homo-multimers, only the latter is used.

#### Multimer structural model ranking

MULTICOM applies three model ranking methods to rank the multimer models. Firstly, the average pairwise structural similarity (PSS) score between a model and other models in the model pool of a multimer is used to rank the structural models^29^. The pairwise structural similarity score is calculated by MM-align^30^. Secondly, the confidence score generated by AlphaFold-Multimer for each model is also used to rank the models. Finally, the average of the two is applied to rank the models.

#### Foldseek structure alignment-based model refinement (FSAMR)

Given an initial multimer model and its MSAs (i.e., MSA_unpaired_ and/or MSA_paired_), the tertiary structure of each subunit in the multimer structural model is used as input for Foldseek to search for similar structures in the pdb_complex template database and the AlphaFoldDB (the version 1 released before March 2022). The structure alignments between each subunit and structural hits are converted into sequence alignments. The sequence alignments of the subunits generated from the Foldseek search are concatenated if they are from the same PDB complex structure or the non-overlapped regions of the same single-chain AlphaFoldDB model to construct the MSA for the multimer. The top structural hits of the subunits are concatenated in a similar way to be used as the templates for the multimer. The concatenated MSAs are added to the original MSA_paired_ to generate a *deeper* MSA. The augmented MSA_paired_, original MSA_unpaired_ (if any for hetero-multimers) and the templates are used as inputs for the customized AlphaFold2-Multimer to generate the refined models. If the highest confidence score of the newly refined models is higher than that of the input model, the refinement process is repeated with the refined model and its MSAs as input until the number of the refinement iterations reaches 5. The refined model with the highest confidence score generated in the refinement process is used as the final output model.

### 2.2 Implementation of the CASP15 assembly structure predictors

During CASP15, the MULTICOM protein assembly structure prediction system was mainly executed on three NVIDIA A100 GPUs with the memory of 40GB, 40GB and 80GB respectively to generate the models for multimer targets before the server prediction deadline and additional models for some multimer targets between the server prediction deadline and the human prediction deadline if necessary. Generally, about 10 - 195 models were generated for each target, depending on its size. The two CASP15 multimer server predictors (MULTICOM_qa and MULTICOM_deep) mainly used the AlphaFold2-Multimer confidence score and the average of the confidence score and the PSS score to rank multimer models, respectively.

The two human multimer predictors (MULTICOM and MULTICOM_human) considered all the models generated before the human prediction deadline. Moreover, the Foldseek structure alignment-based multimer model refinement was applied to refine the top ranked models of most targets and the refined models were added to the model pool for the final multimer model ranking. On average, about 120 models were generated for each human target. MULTICOM_human mainly used the average of the confidence score and the PSS score to rank and select models for final submission, while MULTICOM mainly applied the PSS score to rank models. The ranking may be manually adjusted according to human inspection.

For some very large complexes (e.g., H1111, H1114, H1135, H1137, T1115o, T1176o and T1192o), no full-length multimer models or only poor full-length models could be generated by AlphaFold-Multimer due to the GPU memory limitation, the template-based structure modeling based on Modeller^31^ was applied to combine the models of the components of the complexes generated by AlphaFold-Multimer.

## 3 Results

### 3.1 The comparison between MULTICOM servers and other CASP15 server predictors

According to the CASP15 official assessment (see the official ranking https://predictioncenter.org/casp15/zscores_multimer.cgi), MULTICOM_qa and MULTICOM_deep servers ranked 3^rd^ and 5^th^ among all CASP15 assembly server predictors. The MULTICOM human predictors (MULTICOM_human and MULTICOM) ranked 7^th^ and 10^th^ among all CASP15 assembly predictors. The official CASP15 ranking metric (https://predictioncenter.org/casp15/doc/presentations/Day2/Assessment_Assembly-CASP_EKaraca.pdf) to score a model in a pool of models for a target is 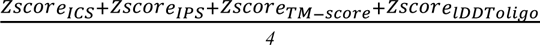, which is the average of Z-scores of ICS (Interface Contact Score)^32^, IPS (Interface Patch Score)^32^, TM-score calculated by US-align^33^ and lDDToligo (Oligomeric lDDT)^34^. Such a score was calculated for the no. 1 model for each target submitted by each predictor. The sum of all positive Z-scores for all the CASP15 targets is the total score of a predictor, which is used to rank all the predictors as shown in **Table 2**. In addition to the top 1 submitted models, CASP15 also calculated the total score for the best of five models for the targets submitted by a predictor to rank the predictors alternatively. According to the Z-scores in terms of the best of five models, MULTICOM_deep, MULTICOM_qa ranked 2^nd^ and 3^rd^ among 26 server predictors. MULTICOM_human and MULTICOM ranked 10^th^ and 14^th^ among 84 predictors.

**Table 2.** The top 15 out of 26 server predictors including NBIS-AF2-multimer ranked by the CASP15 official Z-score and the average TM-score of top 1 models predicted by them for the 41 multimers, 14 TBM multimers, 27 TBM/FM and FM multimers. When calculating the average TM-score here, if a predictor did not submit a prediction for a target, the TM-score for the target is set to 0. The bold font highlights the best result. The underline denotes the second best result.

| Server predictors | Sum of Z-scores (> 0.0) | Target count | Avg TM-score on 41 multimers | Avg TM-score on 14 TBM multimers | Avg TM-score on 27 FM and FM/TBM multimers |
| --- | --- | --- | --- | --- | --- |
| Yang-Multimer<sup>35</sup> | **24.6946** | 38 | 0.7138 | 0.8235 | 0.6569 |
| Manifold-E<sup>36</sup> | <u>18.8589</u> | 41 | **0.7665** | 0.8211 | **0.7382** |
| MULTICOM_qa<sup>37</sup> | 18.3529 | 41 | <u>0.7565</u> | 0.8111 | <u>0.7281</u> |
| DFolding-server<sup>38</sup> | 17.0135 | 33 | 0.5978 | 0.6634 | 0.5637 |
| MULTICOM_deep<sup>37</sup> | 16.2869 | 41 | 0.7416 | **0.8459** | 0.6875 |
| UltraFold_Server<sup>39</sup> | 15.7081 | 41 | 0.6961 | 0.7884 | 0.6483 |
| MultiFOLD<sup>40</sup> | 15.2358 | 41 | 0.6643 | 0.7366 | 0.6268 |
| MUFold<sup>41</sup> | 14.0905 | 41 | 0.7195 | <u>0.8401</u> | 0.6569 |
| Kiharalab_Server<sup>42</sup> | 13.5184 | 40 | 0.6703 | 0.7597 | 0.624 |
| ColabFold<sup>43</sup> | 12.7694 | 39 | 0.6339 | 0.7148 | 0.592 |
| NBIS-AF2-multimer<sup>44</sup> | 12.271 | 41 | 0.7186 | 0.8163 | 0.668 |
| RaptorX-Multimer<sup>45</sup> | 11.9178 | 40 | 0.6744 | 0.78 | 0.6196 |
| Yang-Server<sup>35</sup> | 10.495 | 19 | 0.3578 | 0.6174 | 0.2232 |
| DFolding-refine<sup>38</sup> | 9.3295 | 36 | 0.6584 | 0.8144 | 0.5775 |
| GuijunLab-Assembly<sup>46</sup> | 8.6 | 41 | 0.6701 | 0.769 | 0.6188 |

The average TM-scores of top 1 models (or best of five models) predicted for 41 multimer targets by the top 15 CASP15 server predictors including the standard AlphaFold-Multimer (i.e., NBIS-AF2-multimer run by the Elofsson Group) according to the CASP15 official Z-score ranking are reported in **Table 2** (or **Table S3**). The TM-score of a model is calculated by using US-align^33^ with parameters (*-TMscore 6 -ter 1*) to compare it with the native structure. 41 multimeric targets include 20 hetero-multimers and 21 homo-multimers. A target is classified as template-based modeling (TBM) target if a rather complete template could be found for it and its subunits, while a target was classified as free-modeling (FM) or FM/TBM target if no template or only partial template could be found for it or its subunits. Out of 41 multimeric targets, 14 of them are classified as TBM targets, 27 of them are classified as FM or FM/TBM targets.

The average TM-score of top 1 models submitted by our best MULTICOM server predictor (MULTICOM_qa) for the 41 multimers is 0.7565, only slightly lower than the highest score 0.7665 of the no. 2 Z-score ranked server predictor Manifold-E. It is worth noting that CASP15 Z-score based ranking is not the same as the average TM-score based ranking because the former favors the predictors that perform well on some targets when most other targets fail, which is different from the latter weights all the targets equally. This is the reason Yang-Multimer ranked no. 1 in terms of Z-score even though it missed three targets. The average TM-score of top 1 models of MULTICOM_qa is 5.27% higher than 0.7186 of NBIS-AF2-multimer, showing that a pronounced improvement has been made over the standard AlphaFold-Multimer. Like all the other server predictors, MULTICOM_qa and MULTICOM_deep performed better on the TBM targets than on the FM and FM/TBM targets. MULTICOM_deep has the highest average TM-score of 0.8459 on the TBM targets. MULTICOM_qa has the second highest average TM-score of 0.7281 on the 27 FM and FM/TBM targets, only lower than 0.7382 of Manifold-E.

MULTICOM_qa performed obviously worse than the top-ranked server predictors – either Manifold-E or Yang-Multimer - (i.e., TM-score difference > 0.08) on three homo-trimers (T1174o, T1179o and T1181o), two large hetero-multimers (H1114 and H1137), three hetero-dimers including two nanobodies (H1141 and H1144) and an antibody-antigen H1129, and a large homo-multimers (T1176o) (see **Figure 3**). For T1174o, T1179o and T1181o, MULTICOM_qa managed to generate some good models in the model pool, but the ranking method failed to select them as top 1 model. For H1114 (stoichiometry: A4B8C8), because there was no sufficient GPU memory for AlphaFold-Multimer to generate full-length models for the entire complex of 7,988 residues, MULTICOM_qa tried to build models for different components of the complex (e.g., ABC, AB2C2, A3B3C3, AB4C2, B8) and then combined them to build the structure for the entire complex. Unfortunately, it did not try to build the structure of the A4 component, which is the key to link all the components together. Therefore, MULTICOM_qa submitted a structure predicted for AB2C2 as the top 1 mode as shown in **Figure 3**.

**Figure 3.**
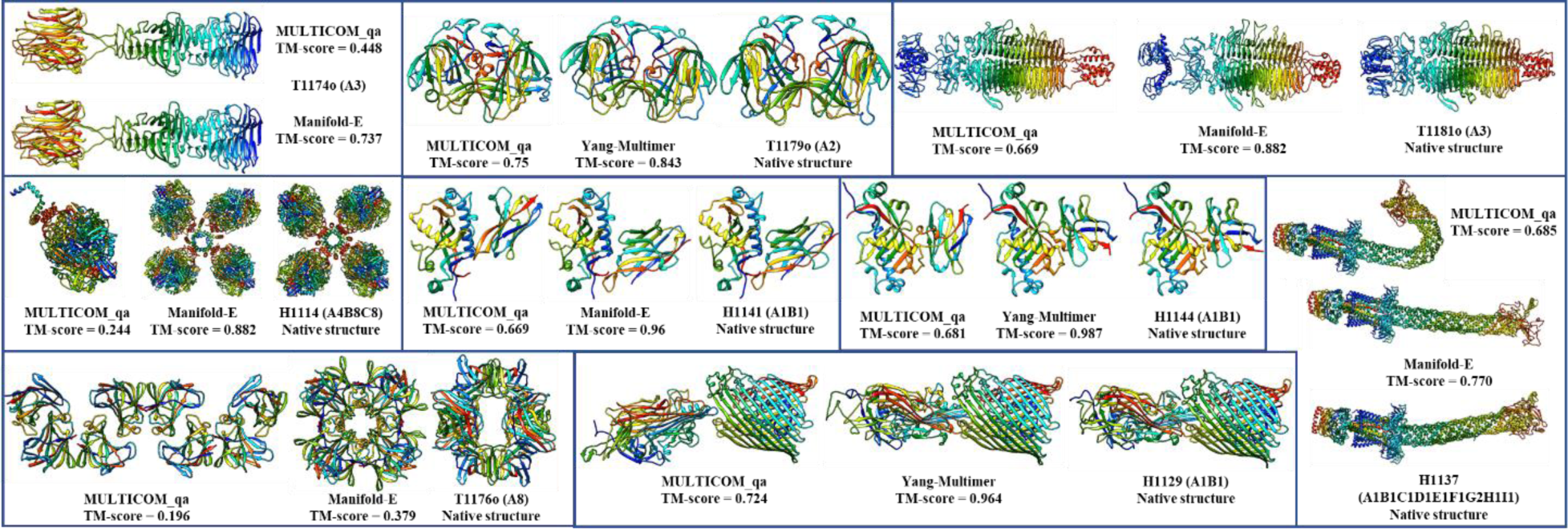
The top 1 submitted models for 9 targets on which MULTICOM_qa performed worse than Manifold-E or Yang-Multimer. The native structure of T1174o is not shown due to the restriction.

For H1137 (stoichiometry: A1B1C1D1E1F1G2H1I1), MULTICOM_qa predicted two conformations for the six-chain transmembrane helical channel consisting of Chains A, B, C, D, E and F (one relatively straight one and one bended one), but it selected the less accurate bended one according to the pairwise similarity between models, leading to the mediocre quality of predicted structures for the target (see **Figure 3**).

For two nanobody-antigen complexes H1141 and H1144 (stoichiometry: A1B1), there are different reasons for the failure. For H1141, the maximum TM-score of the models generated by MULTICOM is 0.6838, much lower than 0.96 of the top 1 model submitted by Manifold-E. For H1144, although some good structural models (TM-score = ∼0.89) had been generated, the ranking method selected a common conformation of low quality rather than the high-quality models that was rare in the model pool. For nanobody targets like H1141 and H1144, it would be useful to generate a large number of models to obtain more high-accuracy models that may have obviously higher confidence scores than other low-quality models. Moreover, as nanobody targets do not necessarily have inter-chain co-evolution information recorded in their MSAs, it may be useful not to pair their MSAs when using AlphaFold-Multimer to generate models for them as shown by some predictors in CASP15.

For H1129, our multimer alignment pairing method did not find any pairs for the two subunits. Therefore, only several default MSAs and templates combinations (e.g., *default_multimer*, *default_pdb*, *default_pdb70*, *default_comp*, *default_struct*, *default_af*) were used as inputs for AlphaFold-Multimer to generate 30 models for it. The maximum TM-score of the generated models is 0.8149, much lower than 0.964 of the top 1 model submitted by Yang-Multimer. T1176o is a homo-multimer with 8 subunits that interact via multiple interfaces, for which AlphaFold-Multimer could generate full-length complex structures directly. However, the interaction between the 8 subunits could not be well predicted by AlphaFold-Multimer. Predicting the structure for 2 or 3 units (A2 or A3) of the multimer produced several different conformations and interfaces, which could not be easily combined to generate good full-length models for the complex. In fact, no model predicted by all the CASP15 predictors have TM-score > 0.5, indicating this is a very hard target.

### 3.2 Overall performance of MULTICOM_qa compared with the standard AlphaFold-Multimer

**Figure 4** shows the distribution of TM-scores of the best of five models submitted by MULTICOM_qa on the 41 multimers (14 TBM multimers and 27 TBM/FM and FM multimers). For 31 out of 41 (75.61%) targets, it generated at least one model with TM-score >= 0.7. For 24 out of 41 (58.54%) targets, it generated at least one high-quality model with TM-score > 0.85.

**Figure 4.**
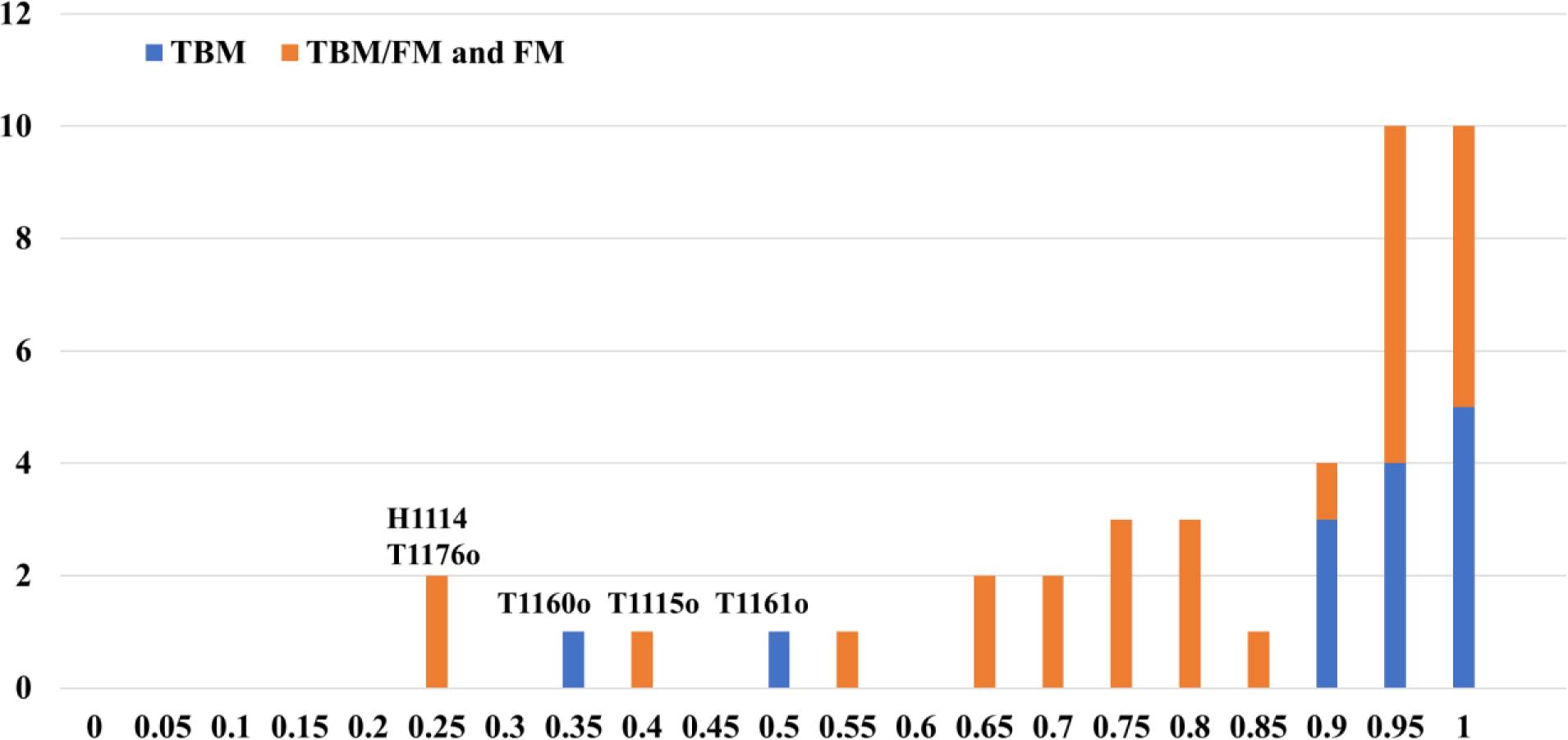
The histogram of the best TM-scores of the top 5 models submitted by MULTICOM_qa on the 27 TBM/FM and FM targets, and 14 TBM targets. The per-target average TM-score of the best model is 0.7963 on the 41 multimer targets.

However, MULTICOM_qa performed very poorly on H1114, T1176o, T1115o, T1160o and T1161o (TM-score of the best model < 0.5). H1114 (stoichiometry: A4B8C8), T1176o (stoichiometry: A8) and T1115o (stoichiometry: A16) are very large multimers. The reason why MULTICOM_qa failed on H1114 and T1176o is explained in Section 3.1. T1115o is a large homomultimer with 16 subunits (4608 residues), for which AlphaFold-Multimer could not generate full-length complex structures due to the lack of GPU memory. Therefore, MULTICOM_qa tried to build the models for A4 and A8, which were combined into an arc-like structure for the complex whose bending angles were different from the native structure.

The two homodimers (T1160o and T1161o) have two very short chains (48 residues only). They have very similar sequences (only five-residue difference in the sequence of the chain) but fold into two different conformations due to different crystallization conditions, which may make it harder for AlphaFold-Multimer to predict their structures. In fact, few CASP15 predictors made good predictions for these two targets, even though AlphaFold-Multimer assigned very high confidence scores (e.g., > 0.8) to the incorrect models that it predicted, indicating they may be outliers.

**Figure 5** compares the TM-score of the best of top 5 models that MULTICOM_qa predicted for each of 41 multimers against that predicted by the standard AlphaFold-Multimer - NBIS-AF2-multimer. On almost all the targets, MULTICOM_qa was able to generate models with quality better than or similar to NBIS-AF2-multimer. Particularly, MULTICOM_qa performed substantially better than NBIS-AF2-multimer on nine targets (H1111, T1187o, T1173o, H1135, H1137, T1179o, T1181o, T1123o and T1115o).

**Figure 5.**
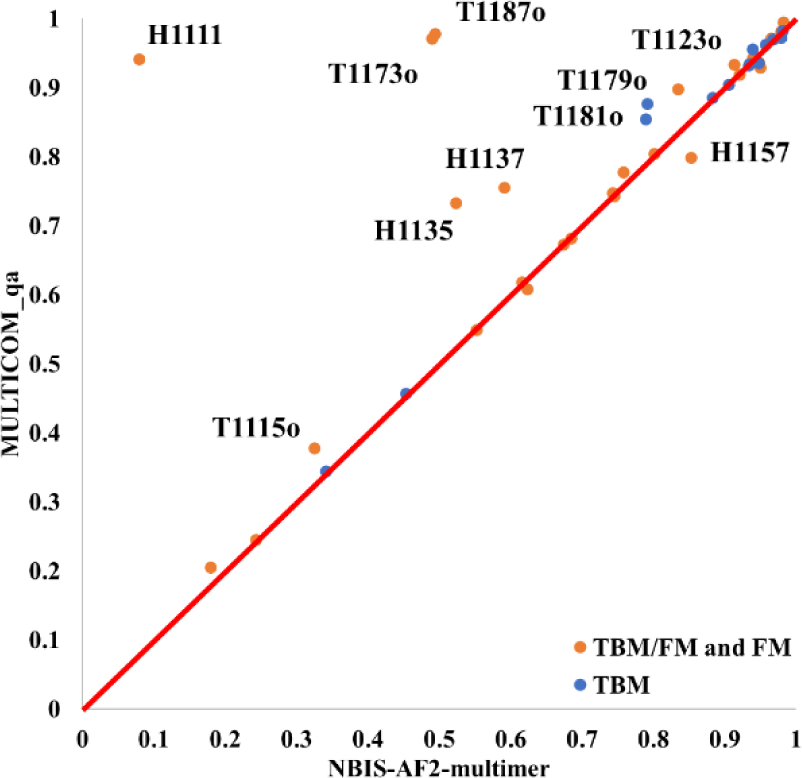
The plot of the TM-score of the best of the top 5 models predicted by MULTICOM_qa for each target against that of NBIS-AF2-multimer on 41 multimer targets.

For nine targets (H1111, T1187o, T1173o, H1135, H1137, T1179o, T1181o, T1123o and T1115o), MULTICOM_qa has an obviously higher TM-score (e.g., difference > 0.05) than NBIS-AF2-multimer, while NBIS-AF2-multimer only has an obviously higher score (e.g., difference > 0.05) than MULTICOM_qa for only one target (H1157). The average best TM-score of the top 5 models of MULTICOM_qa on the 41 multimer targets is 0.7963, which is about 8.0% higher than 0.7375 of NBIS-AF2-multimer. The p-value of the difference is 0.026 according to one-sided Wilcoxon signed rank test. **Figure 6** illustrates nine examples (H1111, T1187o, T1173o, T1115o, H1135, T1181o, T1123o, T1179o, H1137) on which MULTICOM_qa substantially outperformed NBIS-AF2-multimer.

**Figure 6.**
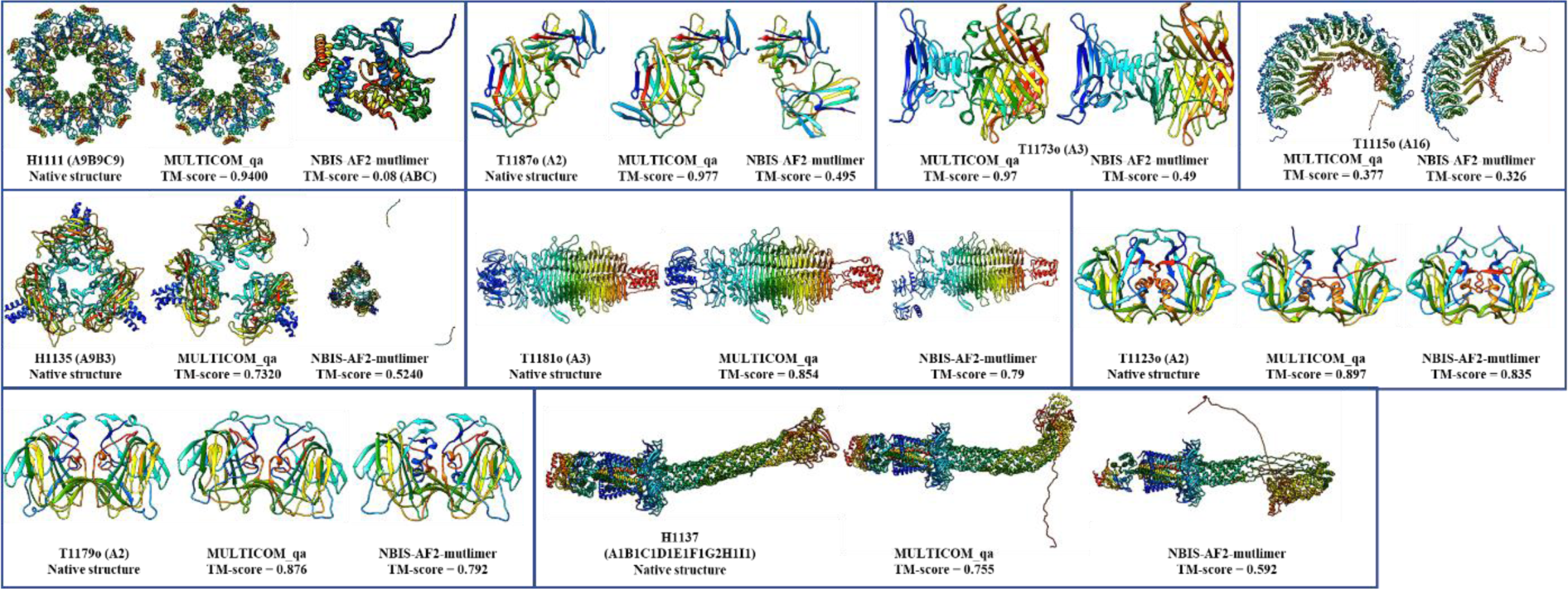
The nine examples (H1111, T1187o, T1173o (native structure not shown due to restriction), T1115o (native structure not shown due to restriction), H1135, T1181o, T1123o, T1179o, H1137) on which MULTICOM_qa performed substantially better than NBIS-AF2-multimer.

### 3.3 Sampling models with diverse multiple sequence alignments and templates improves assembly structure prediction

We compare the best model generated from the MSA-template combinations in **Table S1** and **Table S2** with that of NBIS-AF2-multimer on each of 31 out of 41 CASP15 assembly targets. 10 targets are not included into this analysis for several reasons: unavailability of native structures for T1115o, T1192o and H1185, multiple structural conformations for H1171 and H1172, and no or few full-length structures (i.e, < 5 models) generated for H1111, H1114, H1135, H1137 and T1176o directly by the customized AlphaFold-Multimer in the MULTICOM server system during CASP15 because there was no sufficient GPU memory. The top 5 models generated by the MULTICOM server system for the 31 targets are selected according to their AlphaFold-Multimer confidence scores.

**Figure 7** compares the TM-score of the best of the five models predicted from the diverse MSAs and templates generated by the sequence alignment component in the MULTICOM server system against that of NBIS-AF2-multimer on the 31 multimer targets. The average TM-score of the best models generated by the MULTICOM server system is 0.813, higher than 0.789 of NBIS-AF2-multimer. The results demonstrate using diverse MSAs and templates generated by different sequence alignment approaches as input for AlphaFold-Multimer to generate more models can improve the quality of the best possible models over the standard AlphaFold-Multimer.

**Figure 7.**
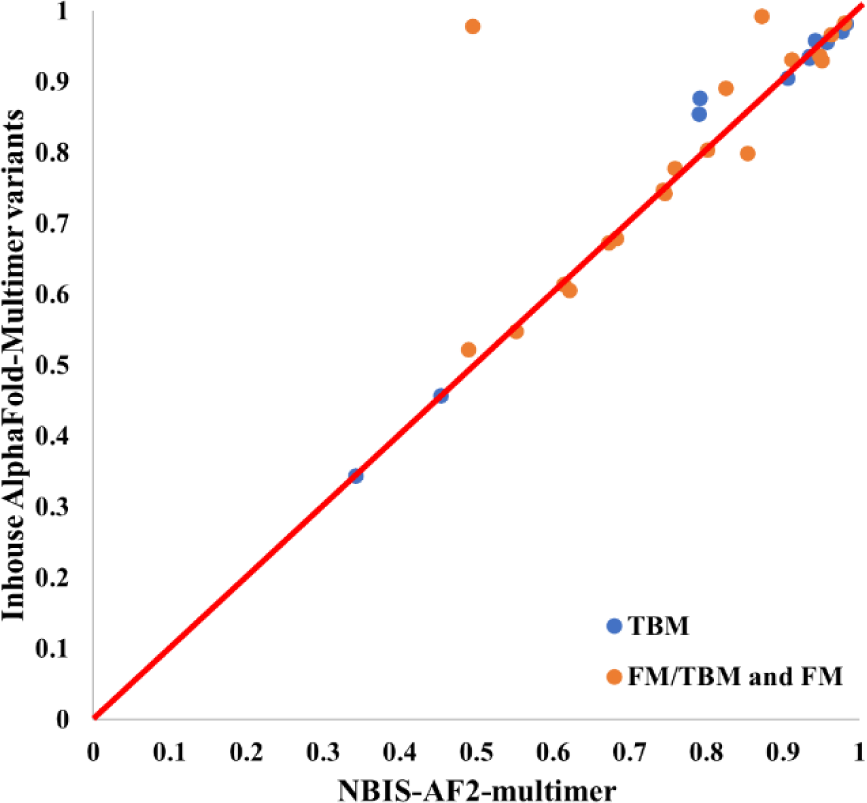
The TM-score of the best of top five models predicted from the diverse MSAs and templates generated by the sequence alignment component in the MULTICOM server system for each of 31 multimer targets (y-axis) against that of NBIS-AF2-multimer (x-axis).

Increasing the value of AlphaFold-Multimer parameter *num_ensemble_eva* from 1 to 3 and *num_recycle* from 3 to 5 and updating the sequence and template databases to the time slightly prior to the start date of CASP15 can also slightly improve the quality of the models generated. For instance, the average per-target best TM-score of using AlphaFold-Multimer with the updated databases and adjusted parameters is 0.8013, slightly higher than 0.789 of NBIS-AF2-multimer. It is worth noting that the sequence databases of NBIS-AF2-multimer were also updated to April 2022 and its template database was updated to May 2022.

### 3.4 Foldseek structure alignment-based model generation improves prediction accuracy

During the CASP15 experiment, the Foldseek Structure Alignment-based Model Generation method (FSAMG) was applied to generate structural models for 26 multimers. For each multimer target, FSAMG was run 2-5 times with different tertiary structures predicted for the subunits/chains of the target, leading to 10 - 25 multimer models generated. On the 26 common targets, the average TM-score of the best of top 5 models ranked by AlphaFold-Multimer confidence score and generated by FSAMG is 0.81, higher than 0.79 of NBIS-AF2-multimer, showing that a noticeable improvement has been made by FSAMG over the standard sequence-alignment-based model generation in AlphaFold-Multimer.

Compared to NBIS-AF2-mutlimer, FSAMG generated much better models on H1140 (0.818 vs 0.622), H1144 (0.890 vs 0.683), T1173o (0.973 vs 0.490), and T1123o (0.893 vs 0.825) as shown in **Figure 8** and **Figure 9** due to several factors below, respectively.

**Figure 8.**
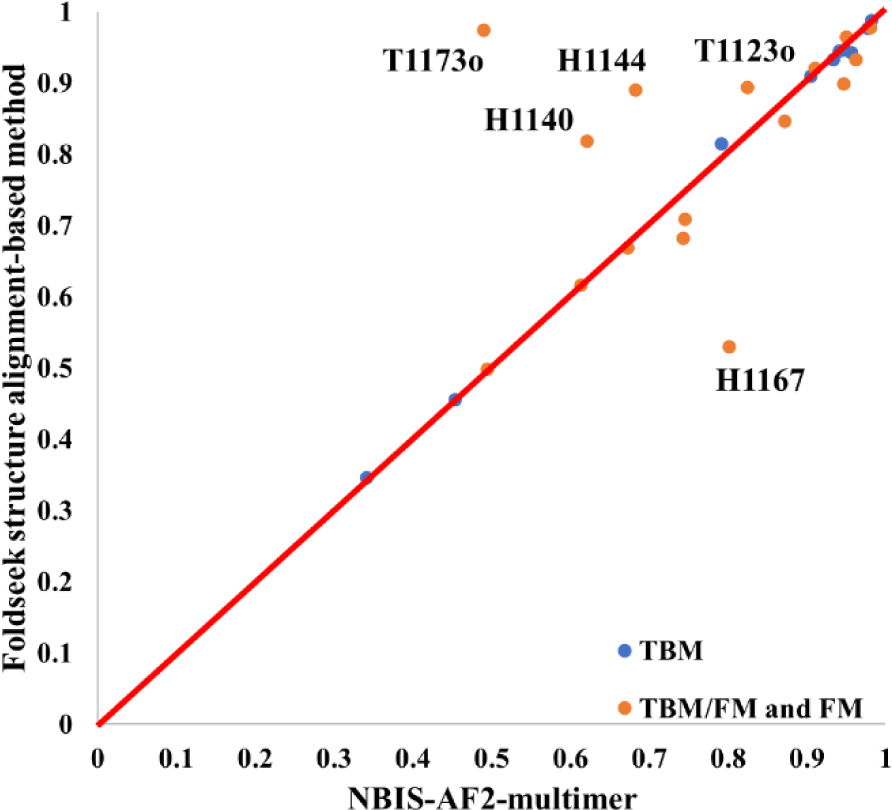
The TM-score of the best of top 5 models for each target generated by Foldseek structure alignment-based model generation (FSAMG) versus NBIS-AF2-multimer on 26 CASP15 multimer targets.

For H1140 (stoichiometry: A1B1), a nanobody target, the MSA for each subunit/chain found by the sequence search (i.e., MSA_unpaired_) was augmented by structural alignments generated by using Foldseek to search the tertiary structure of each chain against the known structures in the PDB. The augmented MSA_unpaired_ of each chain and the similar structural templates found by the Foldseek search were used as input for AlphaFold-Multimer to generate models, even though no paired alignments covering the two chains of H1140 were found by the Foldseek search. The highest TM-score of the models generated by FSAMG is 0.818 (**Figure 9**), much higher than 0.622 of NBIS-AF2-multimer and 0.626 of our in-house AlphaFold-Multimer with the sequence alignment-based MSAs and templates as input. For H1144 (stoichiometry: A1B1), another nanobody target, the highest TM-score of the models generated by FSAMG is 0.890, higher than 0.683 of NBIS-AF2-multimer and 0.855 of our in-house AlphaFold-Multimer with the sequence alignment-based MSAs and templates as input. The multimer models generated by the two methods have very similar tertiary structures for individual chains. However, the multimer models generated by FSAMG have better interactions between the two chains than NIBS-AF2-multimer. Indeed, the main challenge for this target is to predict the interaction between the two subunits because there is no inter-chain co-evolutionary information in the MSAs of nanobody targets. For FSAMG, AlphaFold-Multimer was provided with the MSA_unpaired_ containing newly added structural alignments as well as only two paired alignments in MSA_paired_ to generate models. In contrast, for both H1140 and H1144, our other AlphaFold-Multimer variants that enabled MSA pairing cannot generate good models with ∼1000 paired alignments in the MSA_paired_. The results indicate using more paired MSAs with AlphaFold-Multimer results in bad predictions for nanobody targets because their two chains do not have co-evolution.

**Figure 9.**
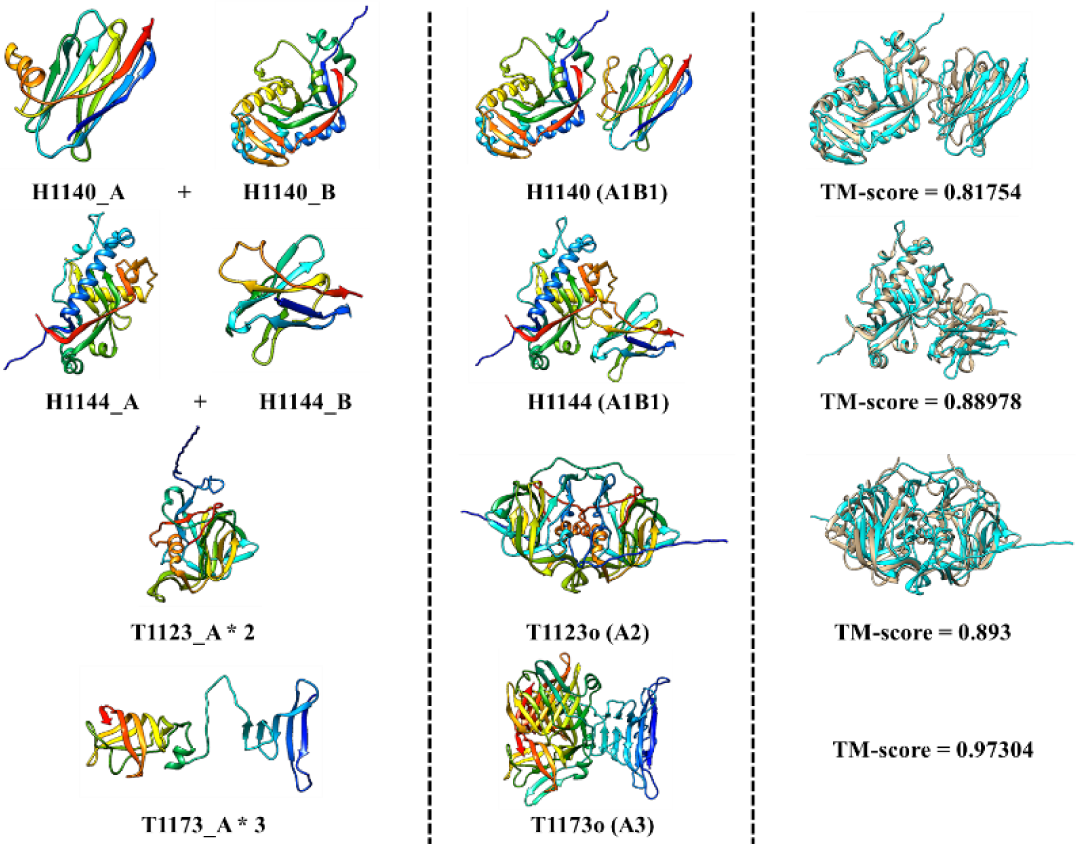
Four good predictions (H1140, H1144, T1123o, and T1173o) made by the Foldseek structure alignment-based multimer model generation (FSAMG). **Left:** input monomer structures generated by AlphaFold2; **Middle**: a multimer model generated by FSAMG; **Right:** the superposition between the multimer model and the native structure (cyan: model, gold: native structure) and the TM-score of the model except for T1173o whose native structure cannot be shown due to the restriction.

For T1123o (stoichiometry: A2), the number of sequences of the initial MSA_paired_ was 26. FSAMG added 10 more structural alignments into the MSA_paired_ and found 4 multimer templates (5W1N, 5KOU, 5KOV, 7RK2) that were fed into AlphaFold-Multimer to generate 10 models. The top 5 models selected by the AlphaFold-Multimer confidence score have TM-scores of 0.873, 0.873, 0.877, 0.893 and 0.881, all higher than 0.825 of NBIS-AF2-multimer.

For T1173o (stoichiometry: A3), the number of sequences in the initial sequence alignment-based MSA_paired_ was already larger than the maximum number of sequences (2048) that can be used by AlphaFold-Multimer, the paired alignments added into the MSA_paired_ by FSAMG made little difference. The main difference is that FSAMG found a new significant multimer template (4UW7) for T1173o that was used as input for AlphaFold-Multimer to generate models. The proportion of high-accuracy models (TM-score > 0.95) generated by FSAMG is 60%, while NIBS-AF2-multimer and our other AlphaFold-Multimer variants did not generate any model of such high accuracy.

Among 26 multimer targets, FSAMG performed only obviously worse than NBIS-AF2-multimer on H1167 - an antibody-antigen target. The best TM-score of its top 5 models is only 0.529, much lower than 0.802 of NBIS-AF2-multimer. The reason is that there were two kinds of conformations (a bad one with TM-score ∼= 0.5 and a good one TM-score ∼= 0.8) in the model pool generated by FSAMG for H1167. The top 5 models selected by the confidence score unfortunately all belong to the bad conformation.

### 3.5. The comparison of the multimer model quality assessment methods

In the CASP15 experiment, the three main quality assessment (QA) methods, including the AlphaFold-Multimer self-reported confidence score (Confidence), the average pairwise similarity score between a model and all other models for a target (PSS) calculated by MM-align, and the average of the two scores (CoPSS), were applied to rank and select multimer models by MULTICOM predictors.

We use the average per-target ranking loss and average per-target correlation to compare the three QA methods on all the full-length models generated for 31 multimers by the CASP15 server prediction deadline (called server_model_dataset) and by the CASP15 human prediction deadline (called human_model_dataset). sever_model_dataset is a subset of human_model_dataset. For some multimer targets, human_model_dataset includes some additional models generated between the server prediction deadline and human prediction deadline. The per-target ranking loss for a target is the difference between the TM-score of the best model for the target in a dataset and the TM-score of the no. 1 model selected for the target by a QA method. Smaller the loss, the better is the ranking for the target. The per-target loss is averaged over all the targets to assess the ranking performance of a QA method. The per-target correlation for a target is Pearson’s correlation between the quality scores for the models predicted by a QA method and the true quality scores (TM-scores) of the models. Higher the per-target correlation, better the predicted quality scores by the QA method. The per-target correlation can be averaged over all the targets to assess the model accuracy estimation (EMA) capability of a QA method.

The average per-target ranking loss and average per-target correlation of the three QA methods on the two datasets are reported in **Table 3**. On the server_model_dataset, CoPSS has the lowest average ranking loss and highest average correlation of 0.0842 and 0.3898, better than 0.0866 and 0.3447 of Confidence and 0.0853 and 0.3767 of PSS, indicating that combining Confidence and PSS improves the performance of estimating the model accuracy of the models in the server_model_dataset. On some targets, PSS can significantly outperform AlplhaFold-Multimer’s confidence score. For instance, PSS’s ranking loss for T1179o is 0.03, much lower than 0.428 of AlphaFold-Multimer’s confidence score.

**Table 3.** The average per-target ranking loss and average per-target correlation of the three QA methods (Confidence, PSS, and CoPSS) on the server_model_dataset and the human_model_dataset.

| QA Method | server_model_dataset |  | human_model_dataset |  |
| --- | --- | --- | --- | --- |
|  | Loss↓ | Correlation↑ | Loss↓ | Correlation↑ |
| Confidence | 0.0866 | 0.3447 | **0.0505** | 0.3845 |
| PSS | 0.0853 | 0.3767 | 0.0892 | 0.3853 |
| CoPSS | **0.0842** | **0.3898** | 0.0625 | **0.4078** |

On the human_server_dataset, Confidence yields the lowest loss of 0.0505 but the lowest correlation of 0.3845, while CoPSS has the second lowest loss of 0.0625 and the highest correlation of 0.4078. Overall, combining Confidence and PSS as CoPSS achieves better performance than PSS. Based on the results on the two datasets, Confidence and PSS are complementary for estimating the accuracy of multimer models. Combining them may be useful to improve the quality assessment of multimer models. However, how to combine them to achieve consistently better results still needs more investigation.

### 3.6. The performance of Foldseek structure alignment-based model refinement

The Foldseek structure alignment-based model refinement method (FSAMR) was applied to 19 multimer targets during the CASP15 experiment. The per-target average maximum TM-scores of the original models is 0.752, similar to 0.750 of the refined models. However, FSAMR was able to generate better models for some targets, especially for T1187o (TM-score 0.899 vs 0.689). For T1187o, the improvement may be due to the extra alignments added to the MSA_unpaired_ by FSAMR. However, FSAMR can also generate models of worse quality. One extreme case is T1153o, where a refined model has a TM-score of 0.484, much lower than 0.928 of the initial model. However, the worse quality can be detected by the change of the AlphaFold-Multimer confidence score of the models. The confidence scores for the 5 original models are close to 0.9, while the confidence scores for the 5 refined models are close to 0.48, indicating a significant drop in the confidence score after the refinement. If we only use the refined models whose confidence score is higher than that of the initial model by at least a margin (i.e., 0.2), the average per-target maximum TM-scores of the refined models is 0.752, higher than 0.740 of the initial models. The results show that FSAMR can be used to generate some diverse and even better models for a multimer target if the change of the model confidence score is substantial.

**Figure 10.**
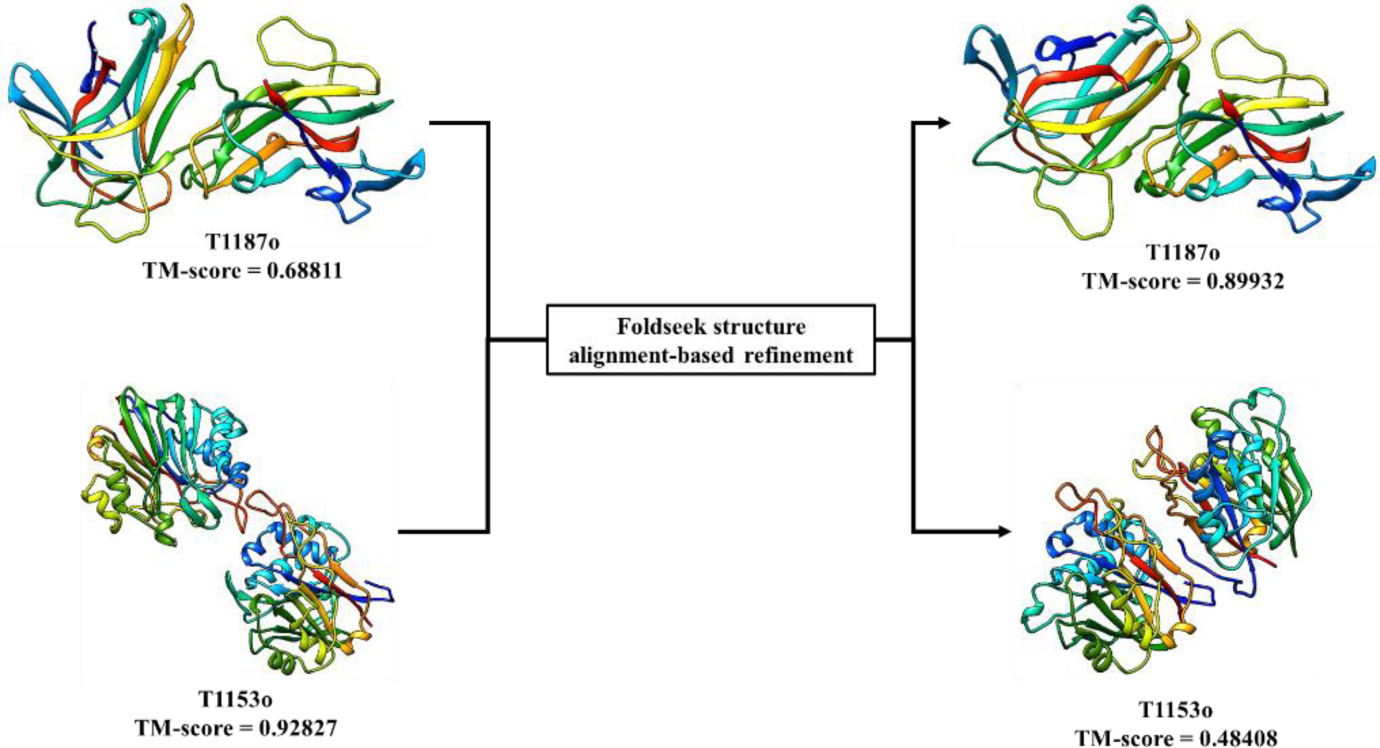
A good example (T1187o) and a bad example (T1153o) for the Foldseek structure alignment-based model refinement (FSAMR). Left: the model before the refinement; Right: the model after the refinement.

### 3.7 MULTICOM server predictors versus human predictors

Compared to the MULTICOM server predictors, the model pool of the MULTICOM human predictors (MULTICOM and MULTICOM_human) was slightly larger since some additional models for some hard targets were generated by either the customized AlphaFold-Multimer with different inputs or by FSAMR between the server prediction deadline and the human prediction deadline. The average TM-score of the best of five models for 41 multimer targets by MULTICOM_qa is 0.796, only slightly lower than 0.797 of the best MULTICOM human predictor (MULTICOM_human), indicating that they achieved largely comparable performance. However, the average TM-score of the top 1 models for the 41 multimer targets of MULTICOM_qa is 0.757, lower than 0.776 of MULTICOM_human. The improvement made by the human predictor comes mostly from the increase of the number of multimer models generated for some targets and some extra human-guided model ranking and combination, especially on the top 1 models. For instance, for T1174o and T1181o, there were two alternative conformations in the top 5 models submitted MULTICOM_qa, but it used the bad conformation as the top 1 model, while MULTICOM_human used the good conformation as top 1 model. For a large hard target T1176o, more structural models were generated by MULTICOM_human for the components of T1176o to be combined to generate full-length models for T1176o. MULTICOM_human’s best model has a TM-score of 0.249, higher than 0.196 of the best model predicted by MULTICOM_qa.

### 3.8 Relationship between multiple sequence alignment (MSA) and multimer model quality

For tertiary structure prediction, the quality of the input MSA quantified by the number of effective sequence (Neff) was shown to have a high correlation coefficient (i.e., 0.777) with the quality score (i.e., GDT-TS) of the tertiary structure models generated by AlphaFold2 for the single-chain monomer targets in CASP15^13^. However, it is more difficult to study the relationship between the quality of MSA and the quality of multimer structural models because AlphaFold-Multimer takes both the MSA of individual chains (MSA_unpaired_) and the paired MSA of the multimer (MSA_paired_) as input, while AlphaFold2 only uses one MSA as input for tertiary structure prediction. Specifically, for homo-multimer consisting of multiple identical chains, AlphaFold-Multimer uses only MSA_paired_ as input, but for hetero-multimer, AlphaFold-Multimer leverages both MSA_unpaired_ and MSA_paired_ if available. Here, we use the results of our default AlphaFold-Multimer variant (*default_multimer* in **Table S1** for homo-multimer and in **Table S2** for hetero-multimer) in the MULTICOM system to study the relationship between MSA quality and model quality. The quality of the input MSAs for a multimer is calculated by Neff_multimer_ 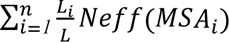, where *n* is the number of subunits of the multimer, MSA_i_ is the combination of MSA_unpaired_ for subunit *i* and the portion of the alignment in MSA_paired_ for subunit *i*, *L_i_*is the sequence length of subunit *i*, *L* is the total sequence length of the multimer. The per-target average correlation between the average TM-scores of the models and Neff_multimer_ of the MSAs is 0.298 on 31 multimer targets, which is a much weaker correlation than the tertiary structure prediction for single-chain monomer targets. The weak correlation may be because the quality of multimer multimers depends not only on the quality of MSAs of individual chains but also the quality of the MSAs informing the interaction between the chains. But this quality of MSAs is not well measured by Neff_multimer_.

### 3.9 Prediction of the structures of very large assemblies

Several multimer targets (e.g., H1111, H1114, H1137 and T1115o) are so large that AlphaFold-Multimer could not generate full-length models for them directly because the 80GB memory of the Nvidia A100 GPU used by MULTICOM was not sufficient to handle them. In this situation, MULTICOM decomposed each of such targets into multiple components to predict the structures of components separately and then combined the structural models of the components into the full-length of the target through the overlapped chains between the components. For instance, H1137 (stoichiometry: A1B1C1D1E1F1G2H1I1) has 9 different chains and 3,939 residues in total. Based on the structure template information, the first domains of six chains (A1B1C1D1E1F1) form a ring, and the ring structure interacts with H and I Chains. Therefore, MULTICOM first predicted the structure of the six chains (A1B1C1D1E1F1) (see supplementary **Figure S1 (A)** for two typical conformations predicted for them: a ring with the straight tail and a ring with the bended tail). It then divided the six chains into a ring structure and a tail structure. The sequences of the ring structure of the first six chains were then cut off to be used with the other three chains (G, H, I) to predict the structure of the 9 chains excluding the tail of the first six chains (see supplementary **Figure S1 (B)**). Finally, the structure of the first six chains and the structure of the 9 chains without the tail were combined by Modeller^31^ through their common ring structure to build the full-length model for H1137 (see supplementary **Figure S1 (C)** for the bended conformation models for H1137 and their TM-score as well as the native structure of H1137). The full-length model with a straight tail (see supplementary **Figure S1 (A)**) has better quality than the one with the bended tail, but the latter is more frequent than the former. The AlphaFold-Multimer confidence score could have selected the structure with the straight tail correctly, but the PSS score preferred the inferior model with the bended tail because it was more abundant.

## 4 Conclusion

We report a new protein prediction system (MULTICOM) to improve AlphaFold-Multimer-based complex structure prediction, which blindly participated in the CASP15 experiment from May to August 2022 as both server and human predictors. MULTICOM enhances AlphaFold-Multimer predictions by generating diverse MSAs and structural templates using both sequence and structure alignments for AlphaFold-Multimer to generate better models, combining AlphaFold-Multimer confidence score with the complementary pairwise model similarity score to rank models, and further refining the models using Foldseek structure alignment to augment MSAs and templates input for AlphaFold-Multimer. MULTICOM_qa server ranked among top CASP15 server predictors for assembly structure prediction and performed significantly better than a standard AlphaFold-Multimer predictor. The results show that using diverse MSAs and structural templates as input is an effective way to generate better models for assembly structure prediction. Particularly, the new Foldseek structure alignment-based model generation (FSAMG) method performs better than the existing sequence alignment-based approach used by AlphaFold-Multimer. Moreover, the Foldseek structure alignment-based model refinement (FSAMR) can substantially improve the quality of structural models for some targets. Furthermore, our results show that the average pairwise similarity between a model and other models is complementary with AlphaFold-Multimer’s self-reported confidence score for estimating the accuracy of assembly models.

## Data Availability

The source code and data of MULTICOM are available at: https://github.com/BioinfoMachineLearning/MULTICOM3.

## Acknowledgements

We thank CASP15 organizers and assessors for making the CASP15 data available. We also thank Dr. Soeding’s group for releasing the latest HHsuite protein hidden Markov model (HMM) database for the community to use prior to the CASP15 experiment. This work is partially supported by two NIH grants [R01GM093123 and R01GM146340], Department of Energy grants [DE-SC0020400 and DE-SC0021303], and three NSF grants [DBI1759934, DBI2308699, and IIS1763246].

## Supplementary Materials

**Table S1.** 19 kinds of combinations of MSA_paired_ and structural templates for homo-multimer targets.

| MSA and Template Combinations | MSA<sub>paired</sub> |  | Structural templates |
| --- | --- | --- | --- |
|  | Sequence database | Interaction source | Template database |
| *default_multimer* | UniRef30, BFD, MGnify clusters | - | pdb70 |
| *default_pdb* | UniRef30, BFD, MGnify clusters | - | pdb_sort90 |
| *default_pdb70* | UniRef30, BFD, MGnify clusters | - | pdb70 |
| *default_comp* | UniRef30, BFD, MGnify clusters | - | pdb_complex |
| *default_struct* | UniRef30, BFD, MGnify clusters | - | pdb_complex |
| *default_af* | UniRef30, BFD, MGnify clusters | - | Predicted chain structures |
| *default_img* | UniRef90, Integrated Microbial Genomes (IMG), metagenome sequence databases | - | pdb70 |
| *uniclust_oxmatch_a3m* | UniClust30 | Species annotation | pdb70 |
| *spec_iter_uniref_a3m* | UniRef30 | Species annotation | pdb70 |
| *spec_iter_uniref_sto* | UniRef90 | Species annotation | pdb70 |
| *spec_iter_uniprot_sto* | UniProt | Species annotation | pdb70 |
| *spec_pdb* | UniRef30 | Species annotation | pdb_sort90 |
| *spec_pdb70* | UniRef30 | Species annotation | pdb70 |
| *spec_comp* | UniRef30 | Species annotation | pdb_complex |
| *spec_struct* | UniRef30 | Species annotation | pdb_complex |
| *spec_af* | UniRef30 | Species annotation | Predicted chain structures |
| *pdb_iter_uniref_a3m* | UniRef30 | PDB | pdb70 |
| *pdb_iter_uniref_sto* | UniRef90 | PDB | pdb70 |
| *pdb_iter_uniprot_sto* | UniProt | PDB | pdb70 |

**Table S2.**
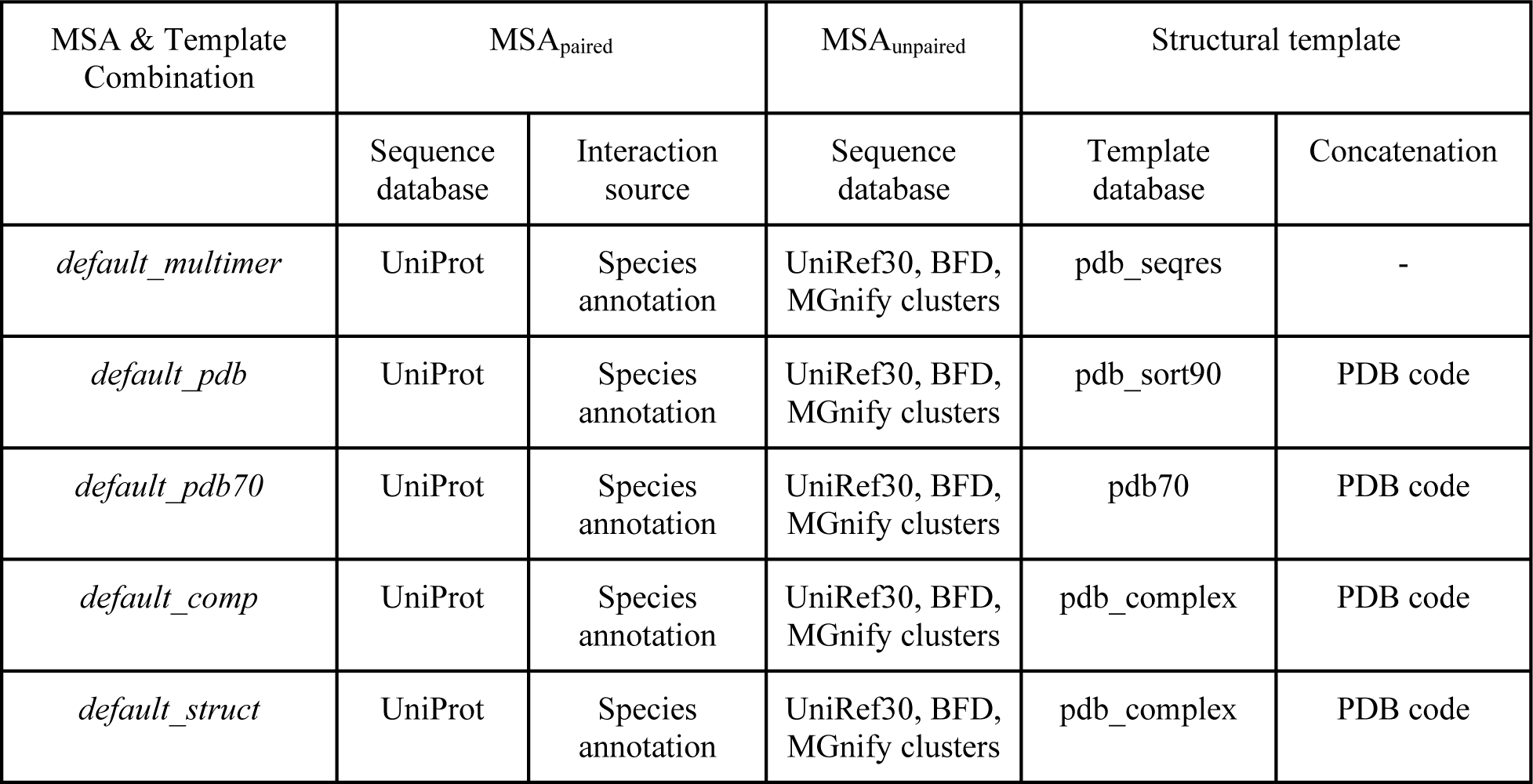
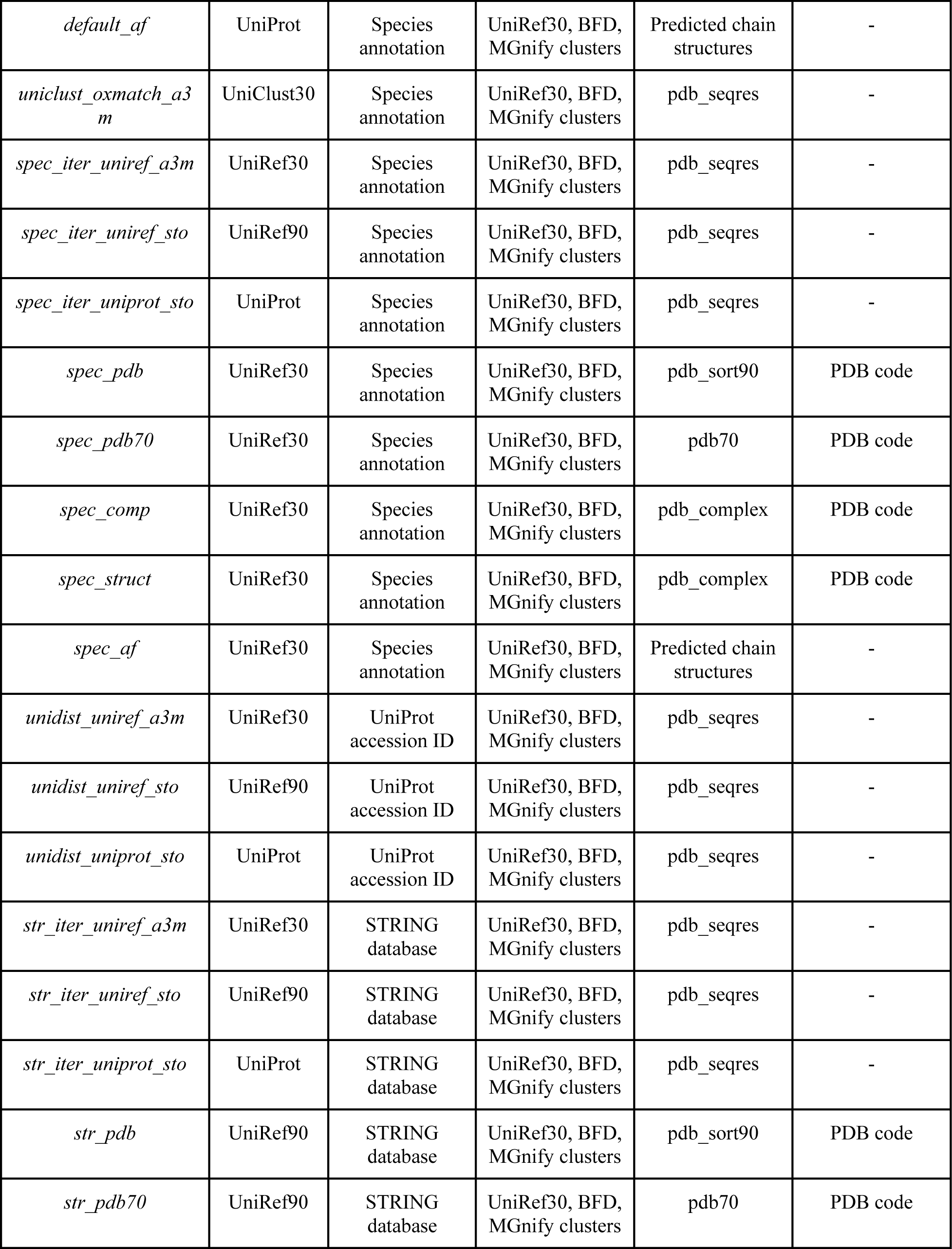
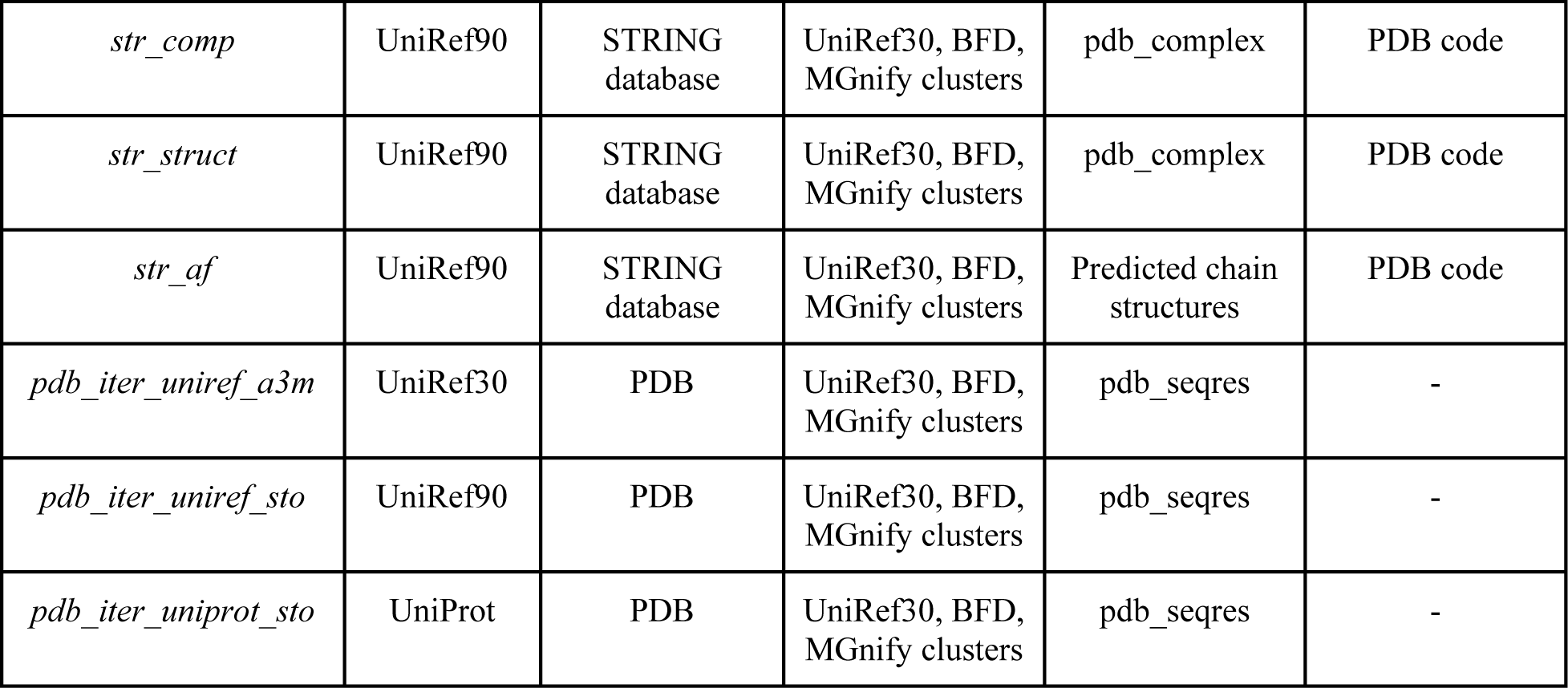
29 kinds of combinations of MSA_paired_, MSA_paired_ and structural templates for hetero-multimer targets.

| MSA & Template Combination | $MSA_{\text{paired}}$ | | $MSA_{\text{unpaired}}$ | Structural template | |
| --- | --- | --- | --- | --- | --- |
|  | Sequence database | Interaction source | Sequence database | Template database | Concatenation |
| *default_multimer* | UniProt | Species annotation | UniRef30, BFD, MGnify clusters | pdb_seqres | - |
| *default_pdb* | UniProt | Species annotation | UniRef30, BFD, MGnify clusters | pdb_sort90 | PDB code |
| *default_pdb70* | UniProt | Species annotation | UniRef30, BFD, MGnify clusters | pdb70 | PDB code |
| *default_comp* | UniProt | Species annotation | UniRef30, BFD, MGnify clusters | pdb_complex | PDB code |
| *default_struct* | UniProt | Species annotation | UniRef30, BFD, MGnify clusters | pdb_complex | PDB code |
| *default_af* | UniProt | Species annotation | UniRef30, BFD, MGnify clusters | Predicted chain structures | - |
| *uniclust_oxmatch_a3m* | UniClust30 | Species annotation | UniRef30, BFD, MGnify clusters | pdb_seqres | - |
| *spec_iter_uniref_a3m* | UniRef30 | Species annotation | UniRef30, BFD, MGnify clusters | pdb_seqres | - |
| *spec_iter_uniref_sto* | UniRef90 | Species annotation | UniRef30, BFD, MGnify clusters | pdb_seqres | - |
| *spec_iter_uniprot_sto* | UniProt | Species annotation | UniRef30, BFD, MGnify clusters | pdb_seqres | - |
| *spec_pdb* | UniRef30 | Species annotation | UniRef30, BFD, MGnify clusters | pdb_sort90 | PDB code |
| *spec_pdb70* | UniRef30 | Species annotation | UniRef30, BFD, MGnify clusters | pdb70 | PDB code |
| *spec_comp* | UniRef30 | Species annotation | UniRef30, BFD, MGnify clusters | pdb_complex | PDB code |
| *spec_struct* | UniRef30 | Species annotation | UniRef30, BFD, MGnify clusters | pdb_complex | PDB code |
| *spec_af* | UniRef30 | Species annotation | UniRef30, BFD, MGnify clusters | Predicted chain structures | - |
| *unidist_uniref_a3m* | UniRef30 | UniProt accession ID | UniRef30, BFD, MGnify clusters | pdb_seqres | - |
| *unidist_uniref_sto* | UniRef90 | UniProt accession ID | UniRef30, BFD, MGnify clusters | pdb_seqres | - |
| *unidist_uniprot_sto* | UniProt | UniProt accession ID | UniRef30, BFD, MGnify clusters | pdb_seqres | - |
| *str_iter_uniref_a3m* | UniRef30 | STRING database | UniRef30, BFD, MGnify clusters | pdb_seqres | - |
| *str_iter_uniref_sto* | UniRef90 | STRING database | UniRef30, BFD, MGnify clusters | pdb_seqres | - |
| *str_iter_uniprot_sto* | UniProt | STRING database | UniRef30, BFD, MGnify clusters | pdb_seqres | - |
| *str_pdb* | UniRef90 | STRING database | UniRef30, BFD, MGnify clusters | pdb_sort90 | PDB code |
| *str_pdb70* | UniRef90 | STRING database | UniRef30, BFD, MGnify clusters | pdb70 | PDB code |
| *str_comp* | UniRef90 | STRING database | UniRef30, BFD, MGnify clusters | pdb_complex | PDB code |
| *str_struct* | UniRef90 | STRING database | UniRef30, BFD, MGnify clusters | pdb_complex | PDB code |
| *str_af* | UniRef90 | STRING database | UniRef30, BFD, MGnify clusters | Predicted chain structures | PDB code |
| *pdb_iter_uniref_a3m* | UniRef30 | PDB | UniRef30, BFD, MGnify clusters | pdb_seqres | - |
| *pdb_iter_uniref_sto* | UniRef90 | PDB | UniRef30, BFD, MGnify clusters | pdb_seqres | - |
| *pdb_iter_uniprot_sto* | UniProt | PDB | UniRef30, BFD, MGnify clusters | pdb_seqres | - |

**Table S3.** The average TM-score of the best of five models of the top 15 out of 26 server predictors including NBIS-AF2-multimer (the standard AlphaFold-Multimer predictor) on the 41 multimers, 14 TBM multimers, 27 TBM/FM and FM multimers. When calculating the average TM-score, if a predictor did not submit a prediction for a target, the TM-score is set to 0. The bold font highlights the best result. The underline denotes the second best result.

| Server predictor | Sum of Z-scores (> 0.0) | Avg TM-score on 41 multimers | Avg TM-score on 14 TBM multimers | Avg TM-score on 27 FM and FM/TBM multimers | Target count |
| --- | --- | --- | --- | --- | --- |
| Yang-Multimer | **28.4019** | 0.7542 | **0.8619** | 0.6984 | 39 |
| MULTICOM_deep | <u>24.6154</u> | 0.7781 | 0.8536 | 0.7389 | 41 |
| MULTICOM_qa | 23.1823 | <u>0.7963</u> | <u>0.8541</u> | <u>0.7663</u> | 41 |
| Manifold-E | 22.8345 | **0.8161** | 0.8504 | **0.7984** | 41 |
| DFolding-server | 20.2187 | 0.6816 | 0.7835 | 0.6287 | 36 |
| ColabFold | 18.3899 | 0.6715 | 0.7584 | 0.6265 | 39 |
| UltraFold_Server | 18.2015 | 0.7641 | 0.7999 | 0.7456 | 41 |
| MultiFOLD | 17.4288 | 0.6900 | 0.7671 | 0.6500 | 41 |
| Kiharalab_Server | 16.561 | 0.6995 | 0.7839 | 0.6557 | 40 |
| MUFold | 16.4772 | 0.7478 | 0.8481 | 0.6958 | 41 |
| NBIS-AF2-multimer | 14.8905 | 0.7375 | 0.8438 | 0.6824 | 41 |
| RaptorX-Multimer | 14.7853 | 0.7061 | 0.8168 | 0.6487 | 40 |
| DFolding-refine | 12.8844 | 0.6973 | 0.8362 | 0.6253 | 36 |
| GuijunLab-DeepDA | 12.6262 | 0.7361 | 0.8309 | 0.6870 | 41 |
| Yang-Server | 12.2828 | 0.3839 | 0.6557 | 0.2430 | 19 |

**Figure S1.**
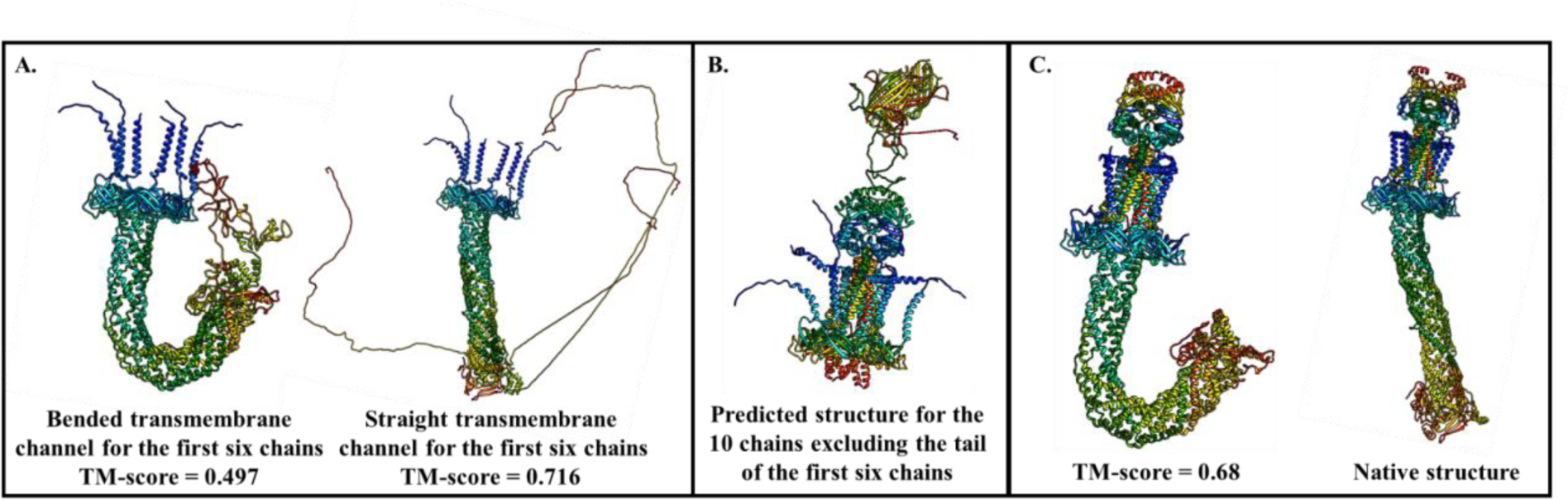
A. The two conformations (with TM-score = 0.497 and 0.716) of predicted complex structure for the first six chains of H1137 (A1B1C1D1E1F1); **B.** The predicted structure for the 10 chains excluding the tail of the first six chains; **C.** The native structure and the full-length complex structure (TM-score = 0.68) for H1137 built from the bended transmembrane channel for the first six chains (**left structure in plot A**) and the predicted structure for the 10 chains excluding the tail of the first six chains (**structure in plot B**).

